# Structural basis for inhibition of βFXIIa by garadacimab

**DOI:** 10.1101/2023.12.03.569829

**Authors:** Ieva Drulyte, Rajesh Ghai, Saw Yen Ow, Eugene A. Kapp, Adam J. Quek, Con Panousis, Michael J. Wilson, Andrew D. Nash, Matthias Pelzing

## Abstract

FXIIa is the principal initiator of the plasma contact system and can activate both procoagulant and proinflammatory pathways. Its activity is important in the pathophysiology of Hereditary Angioedema (HAE). Here, we describe a high-resolution cryo-EM structure of the beta-chain from FXIIa (βFXIIa) complexed with the Fab fragment of garadacimab. Garadacimab binds to βFXIIa through an unusually long CDR-H3 that inserts into the S1 pocket in a non-canonical way. This atypical structural mechanism is likely the primary contributor in the inhibition of activated FXII proteolytic activity in HAE. Garadacimab Fab-βFXIIa structure also reveals critical determinants of high-affinity binding of garadacimab to activated FXII. Structural analysis with other bonafide FXIIa inhibitors, such as benzamidine and C1-INH, reveals a conserved mechanism of βFXIIa inhibition, a novel finding of this study. In summary, garadacimab Fab-βFXIIa structure provides crucial insights into its mechanism of action and delineates primary and auxiliary paratopes/epitopes. This work reaffirms the benefits of cryo-EM as a primary tool to study antigen-antibody complexes in near native state, particularly where other methods have fallen short.

## INTRODUCTION

Hereditary angioedema (HAE) is a rare genetic disorder whereby patients experience recurring episodes of swelling attacks, often accompanied with pain. It is a debilitating and potentially life-threatening disorder that impacts 1 out of 50,000 (Zuraw, 2008). Individuals suffering from this disease often have either low levels (Type I), or normal but dysfunctional levels (Type II), of C1 esterase inhibitor (C1-INH), a critical multifunctional serine protease inhibitor. The current standard of care for HAE involves the use of both on-demand and prophylactic treatments and is achieved through C1INH replacement therapies, kallikrein inhibitors, or bradykinin B2 receptor antagonists. C1-INH is a serpin, a primary endogenous regulator of both the classical complement activation and contact systems (Conway, 2015; Davis et al., 2010), and rapidly inhibits ∼95% of activated FXIIa (de Agostini et al., 1984). FXIIa serves as the central component of procoagulant and proinflammatory plasma contact system. Garadacimab is a novel fully human recombinant monoclonal antibody inhibitor of human activated Factor XII (FXIIa) that has recently completed its Phase 3 trial for the prophylactic treatment of HAE patients with C1-INH deficiency. Results from the most recent clinical trial show that garadacimab was able to significantly reduce HAE attacks in patients aged 12 years and older and had a favourable safety profile (Craig et al., 2023).

Garadacimab is the affinity-matured version of 3F7 which was the parental antibody identified using phage display and is a potent inhibitor of FXIIa and beta-chain of activated factor XII (βFXIIa) (Cao et al., 2015). It has been previously established that the antibody recognizes βFXIIa with high specificity (Larsson et al., 2014). In human plasma, 3F7 is preferentially bound to FXIIa over zymogen (Larsson et al., 2014). The antibody garadacimab binds with high affinity to human βFXIIa (K_D_ = 140 pM), which is a ∼44-fold increment over the parental antibody (Cao et al., 2018). Binding of 3F7 to βFXIIa was enhanced through mutations in the heavy chain complementarity determining region (CDR) 2. 3F7 heavy chain CDR2 residues (GIRPSGGTTVYADSVKG) were mutated to the underlined residues GI<u>DIPTKG</u>TVYADSVKG in the affinity matured garadacimab antibody. The molecular determinants that led to the increase in garadacimab affinity to βFXIIa were unclear. Mutation of D60K/V97K in βFXIIa abolished 3F7 binding. This data provided early insights that 3F7 binding to βFXIIa maps near the 99-loop and 60-helix of βFXIIa. Therefore, it was believed 3F7 functions by blocking the βFXIIa catalytic cleft (Larsson et al., 2014).

Critical to the understanding of the intended pharmacological effect through inhibition of FXIIa are insights into the mechanism of action (MoA) of garadacimab. To this effect, x-ray crystallography was employed to gain a structural understanding of garadacimab Fab in complex with βFXIIa. While reproducible needle-like small crystals of the complex were obtained, the crystals did not diffract. This is potentially due to poor packing in the crystal lattice. Simultaneously, a hydrogen-deuterium exchange mass spectrometry (HDX-MS) study to elucidate the potential binding interface of garadacimab on βFXIIa was initiated. This provided evidence that garadacimab may in fact obstruct entry of the βFXIIa S1 pocket (Ow et al., 2023). Through solvent accessibility data coupled with HDX rate, we could highlight putative residues forming the binding interface. However, a deep understanding of the overall MoA of garadacimab and its biological implications was lacking.

In this investigation, we report a high-resolution cryo-electron microscopy (cryo-EM) structure of βFXIIa in complex with Fab fragment of garadacimab. The structure demonstrates an atypical interaction of CDR-H3 of garadacimab with βFXIIa that inserts deep inside βFXIIa to inhibit proteolytic activity. Structural modelling provided insights into how an endogenous inhibitor (C1-INH) and a therapeutic inhibitor (garadacimab) of FXIIa adopt a conserved mechanism to block access to S1 pocket, resulting in reduced proteolytic cleavage of substrates. Structural analysis also reveals critical determinants of the high-affinity binding of garadacimab to activated FXII. Overall, this study extends and improves our understanding of garadacimab, and its MoA, as well as delineates primary and auxiliary paratopes/epitopes.

## RESULTS

### Structural basis for the inhibition of βFXIIa by garadacimab

To understand the structural basis for garadacimab-mediated inhibition of βFXIIa, we sought to perform cryo-EM analysis on the recombinant human plasma clotting FXIIa in complex with the garadacimab Fab fragment. Initial cryo-EM attempts of the garadacimab Fab-βFXIIa complex resulted in very packed monolayers of particles (**Fig. S3A**) and poor resolution of the resulting reconstructions (data not shown). To solve this issue, we followed the lead of a study utilizing a VHH specific for the human kappa light chain to assist x-ray crystallography efforts (Ereño-Orbea et al., 2018) and added a variable heavy-chain (VHH) domain to the garadacimab Fab-βFXIIa complex, which increased the complex’s mass from ∼77 to ∼90 kDa (**Fig. S1**). Analysis of the ternary complex, concentrated to ∼10 mg/ml for cryo-EM sample preparation, using SEC-MALS revealed a higher molecular weight peak that we hadn’t observed before, estimated to be roughly twice the size of the monomeric complex (**Fig. S2**).

Single particle analysis (SPA) of the data revealed heterogeneity in the conformation of the garadacimab Fab-βFXIIa complex, which aligns with the SEC-MALS results. We observed three distinct garadacimab Fab-βFXIIa complexes: a C2-symmetric dimer (65%), an asymmetric dimer (8%) and an apparent monomer (27%) (**Fig. S4**), which were resolved to 2.6 Å, 3.8 Å and 3.9 Å, respectively (**Fig. S4**). In all cases, the interaction between the βFXIIa and garadacimab Fab was constant (**Fig. 1B, S5I-J**), as such, we decided to use the highest resolution reconstruction for further interpretation of the epitope-paratope interface.

**Figure 1:**
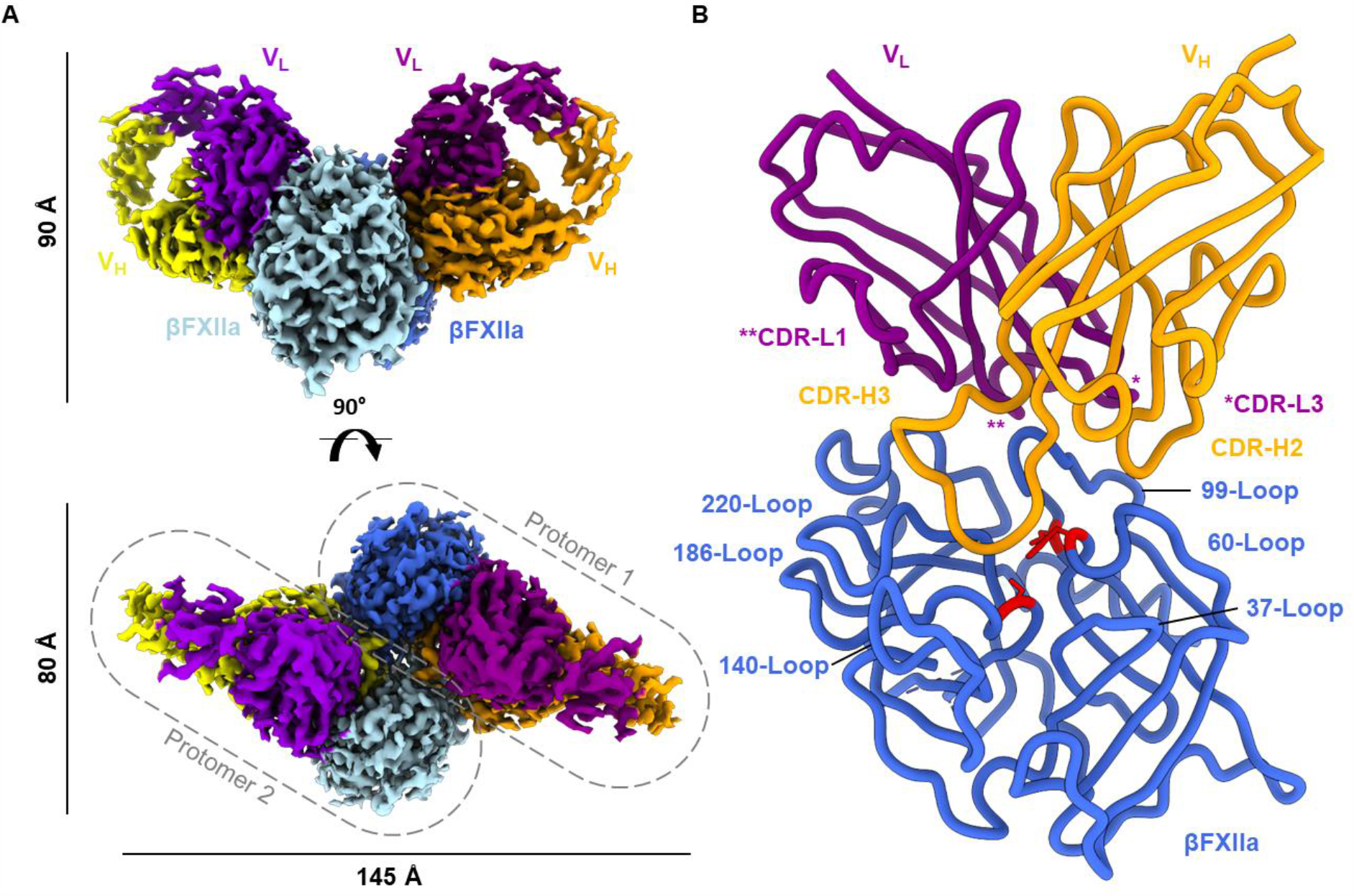
Cryo-EM structure of garadacimab Fab-βFXIIa complex. (A) Surface rendering of the garadacimab Fab-βFXIIa dimer, shown as two orthogonal views. (B) The structural model of garadacimab Fab-βFXIIa complex with βFXIIa, light and heavy chains coloured blue, purple and orange, respectively. Garadacimab CDR loops, βFXIIa surface loops are labelled, and residues of βFXIIa catalytic triad are coloured in red.

Non-uniform refinement of the C2-symmetric dimer produced a density map with a global resolution of 2.6 Å (**Fig. 1A, S3**), with the epitope-paratope region resolved ∼2.4 Å. This resolution permitted the atomic modelling of both the βFXIIa and the Fab variable domains as well as the number of ordered waters involved in the epitope interface (**Fig. 1B**,**2**). Additionally, a density consistent with a bi-antennary core fucosylated glycan was observed at the bottom of the molecule, near the βFXIIa dimerization interface (**Fig. S7A**). Mass spectrometry (Fig. S7B) confirmed the glycan’s identity, and a fucosylated HexNac core with three mannose units was built into the density with high confidence.

Consistent with the previous HDX-MS data (Ow et al., 2023), garadacimab inhibits βFXIIa by binding to the βFXIIa loops neighbouring the S1 catalytic pocket entrance and preventing substrate binding through steric hindrance. The binding of the garadacimab Fab to βFXIIa buries a surface area of ∼1054 Å^2^, with ∼700 Å^2^ from the heavy and ∼355 Å^2^ from the light chain. The paratope is composed of the CDR-H2, H3, L1 and L3 loops, mediated primarily by hydrogen bonds (**Fig. 2**). The resolution of the map was sufficient to visualize several water molecules which participate in the solvent-mediated H-bonding.

**Figure 2:**
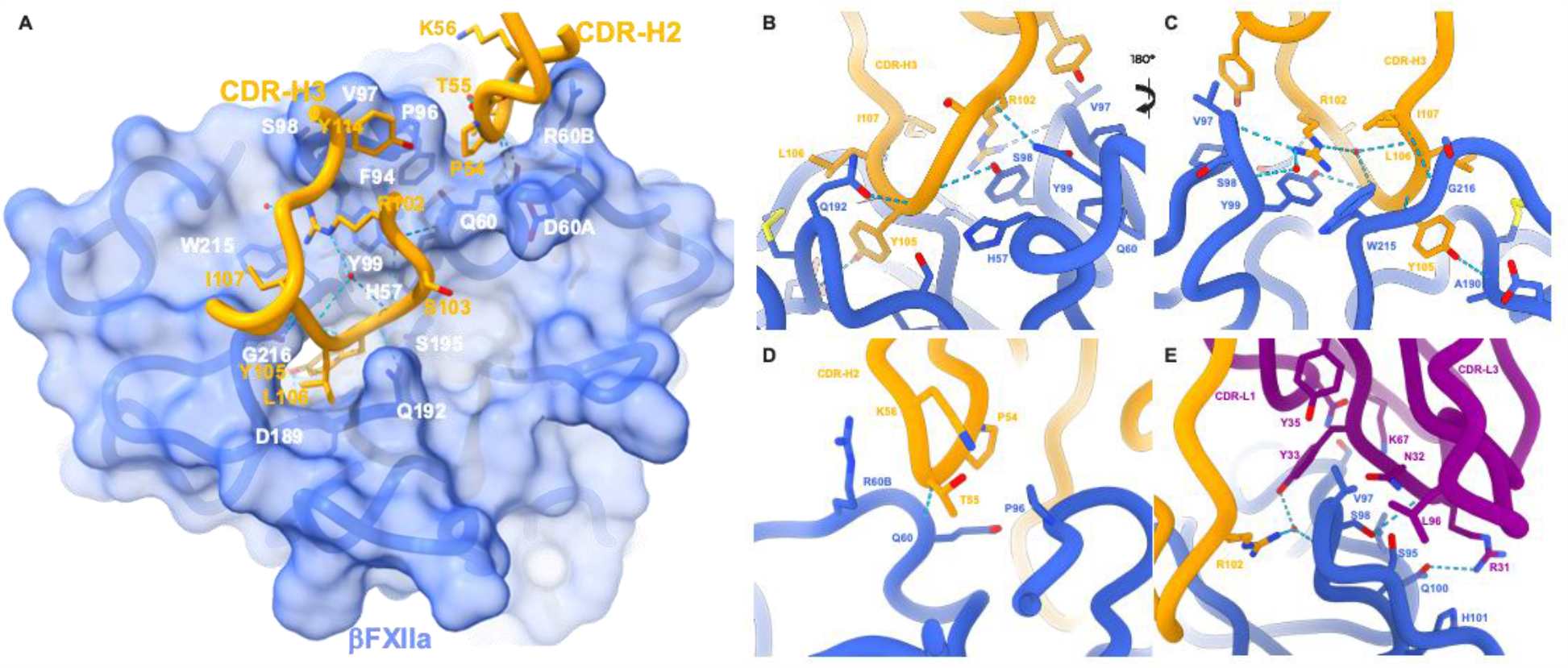
The garadacimab inhibits βFXIIa by binding to the βFXIIa loops neighbouring the S1 catalytic pocket entrance and preventing the substrate binding through steric hindrance. (A) Overall view of a surface representation of βFXIIa (blue) bound to CDR-H3 and CDR-H2 (orange) with side chains of critical contact residues (sticks) responsible for epitope-paratope interactions. (B-E) Close-up view of the garadacimab CDR-H3 (A-B, two orthogonal views), CDR-H2 (C) and CDR-L1 epitopes with the residues involved in the epitope-paratope interactions shown as sticks. Hydrogen bonds are denoted with dashed lines.

A chymotrypsin-based numbering system has been used for consistency with the established literature (Dementiev et al., 2018). Accordingly, CDR-H3 residues Arg102, Tyr105, Leu106 and Ile107 interact with the βFXIIa residues Gln60, Val97, Tyr99, Ala190, Trp215 and Gly216, with Arg102 being involved in the water-coordinated H-bond network with βFXIIa residues Ser98 and Trp215 (Fig. 2A-C. CDR-H2 residue Thr55 H-bonds to Gln60 (Fig. 2D). CDR-L1 and L3 residues Arg31, Asn32, Tyr33 and Leu96 interact with βFXIIa residues Ser95, Ser98 and Gln100 (Fig. 2E. A water-mediated H-bond between CDR-H3 and CDR-L1 also stabilizes the paratope. βFXIIa dimerization interface reveals H-bonds between Gly147-Gln192 and Asp149-Tyr151 with an additional solvent-mediated H-bond to Phe41 (Fig. 3).

**Figure 3:**
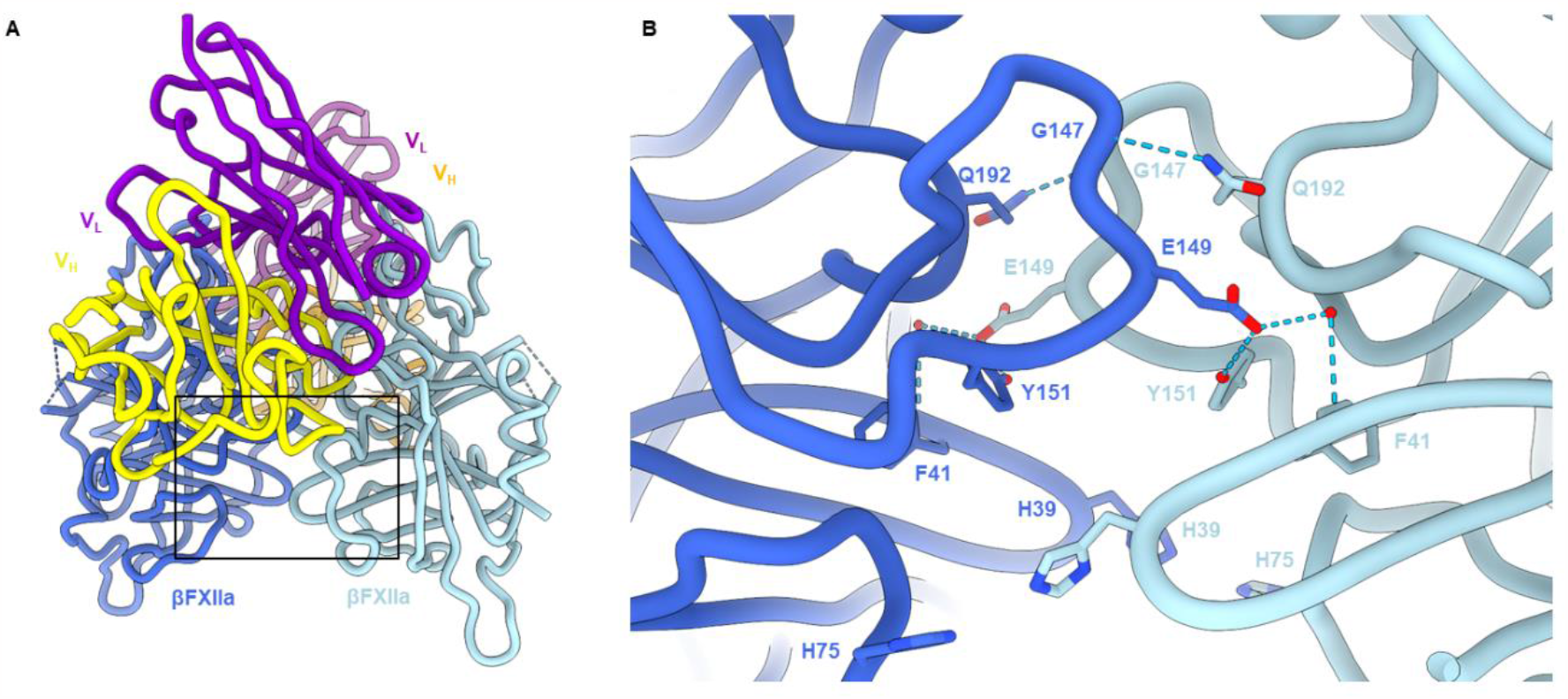
Cryo-EM structure of garadacimab Fab-βFXIIa complex reveals that antigen forms a homodimer and further corroborates HDX-MS solvent accessibility data. (A) Garadacimab Fab-βFXIIa atomic model, with the boxed region representing the βFXIIa dimerization interface. (B) Zoomed-in view of the dimerization interface showing that dimerization is mediated through an interface comprising glutamic acid at position 149 and Tyrosine 151. Fab heavy and light chain models are excluded from this panel for clarity.

### Garadacimab possess a conserved mechanism for βFXIIa inhibition

While benzamidine serves as a non-covalent inhibitor of FXIIa, C1-INH is a primary endogenous covalent inhibitor, inhibiting ∼95% of FXIIa (Dementiev et al., 2018; A de Agostini et al., 1984). In the absence of high-resolution structural information, it is unclear how FXIIa and βFXIIa protease activity gets downregulated by C1-INH. Typically, serpins such as C1-INH engage with the bonafide serpin protease through insertion of the reactive center loop (RCL) into the protease and forming a covalent bond. Atypical CDR-H3 loop insertion into the substrate and inhibitor binding pocket of βFXIIa shares many similarities with C1-INH mode of protease recognition. To study this, we generated a model of the active form of C1-INH in complex with βFXIIa using Alphafold multimer (Evans et al., 2022; Mirdita et al., 2022). Remarkably, the model demonstrates the placement of the RCL into the same binding pocket as CDR-H3 (**Fig. 4A, S8**). P1 residue (Arg444) from the RCL sequence SVARTLLV of C1-INH is known to be a critical determinant of FXIIa activity (Dementiev et al., 2018). Mutagenesis studies have shown that perturbation of Arg444 leads to complete loss of FXIIa inhibition (Eldering et al., 1992). Superposition of garadacimab Fab bound βFXIIa structure, and the benzamidine bound βFXIIa (PDB id: 6b74), on C1-INH bound βFXIIa model revealed complete overlay of guanidine group of Arg444, phenyl ring of Tyr105 and benzene ring of benzamidine. Arg444 from the RCL also mimicked salt bridge formation of Asp189-Tyr105 (garadacimab Fab-βFXIIa complex) and Asp189-aminidium group (benzamidine-βFXIIa complex) (**Fig. 4A**). Moreover, it has been shown that 3F7 competes with C1-INH and blunted complex formation with FXIIa (Larsson et al., 2014). Taken together, these data highlight that garadacimab, C1-INH and benzamidine possess a conserved mechanism for inhibiting FXIIa and βFXIIa.

**Figure 4:**
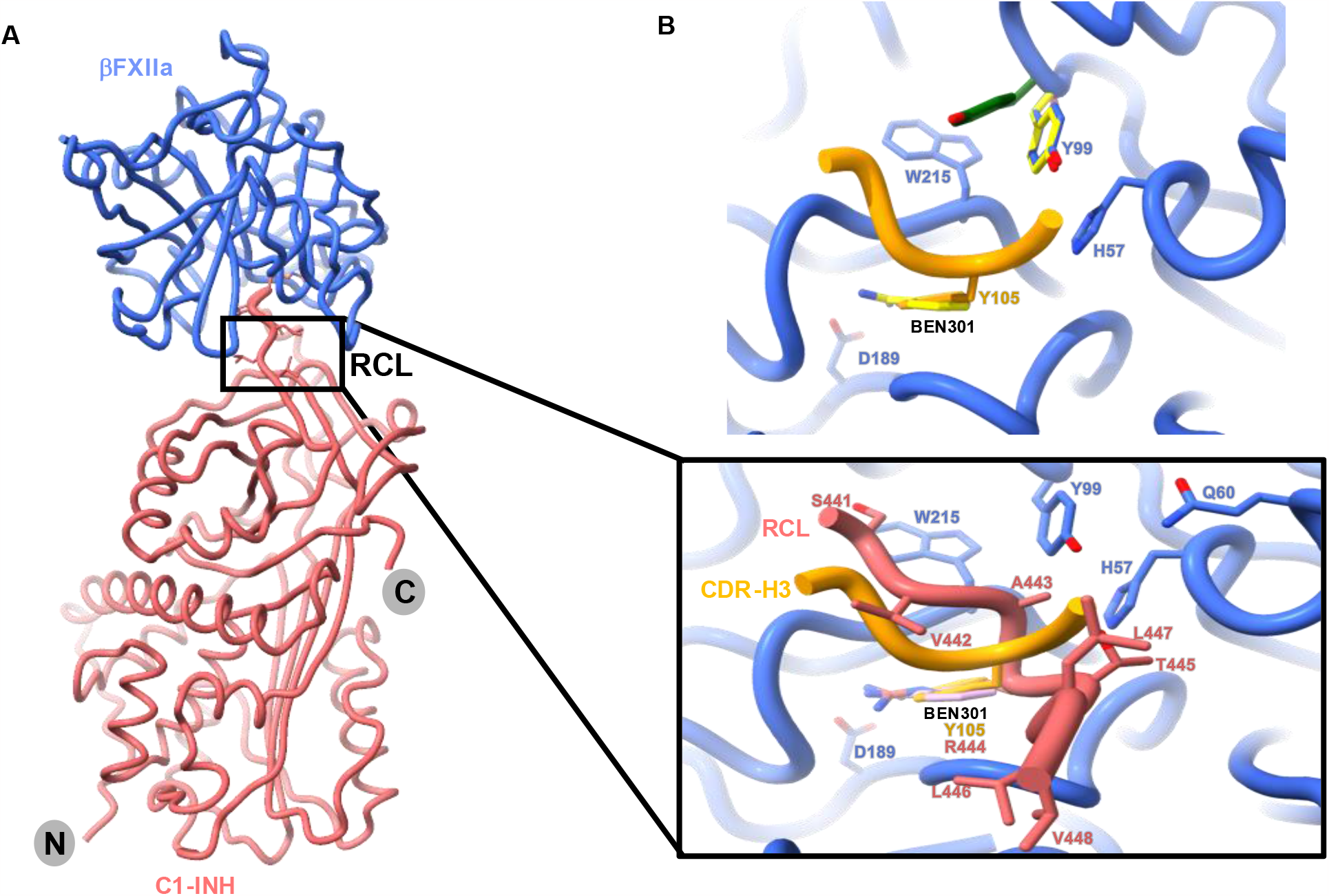
Garadacimab C1-INH and benzamidine possess a conserved mechanism for the inhibition of FXIIa and βFXIIa. (A) Alphafold generated model of βFXIIa (blue)-C1-INH (coral) complex (cartoon). The consensus sequence residues from the RCL are shown as sticks. The inset demonstrates a close-up overlay of garadacimab Fab-βFXIIa structure with the model of βFXIIa-C1-INH complex and structure of benzamidine-bound βFXIIa revealing Tyr105 from CDR-H3 of garadacimab overlays well with the benzamidine ring and critical P_1_ Arg444 from the RCL of C1-INH indicating a conserved inhibition mechanism for βFXIIa. (B) Cartoon demonstrating overlay of garadacimab Fab-βFXIIa in orange and blue, uncomplexed βFXIIa (PDB id: 6gt6) in green and benzamidine bound βFXIIa (PDB id: 6b74) in yellow. A portion of CDR-H3 is shown in orange with the critical Try105 in sticks superimposing well with the benzamidine ring (yellow sticks). Tyr99 from the superposition of garadacimab Fab-βFXIIa (blue sticks), uncomplexed βFXIIa (green sticks) and benzamidine bound βFXIIa (yellow sticks) structure is shown in open and closed forms.

## DISCUSSION

### Structural relevance and Improvements to mechanism of action understanding

Our cryo-EM structure of βFXIIa in complex with garadacimab Fab presents the first example of how βFXIIa function is inhibited. The interaction of Fab with βFXIIa demonstrates an atypical antigen-antibody complex. Typically, the CDRs contact solvent-exposed epitope residues on the surface of the antigen. In this case, garadacimab possesses an unusually long (17 amino acids) CDR-H3 which inserts deep into the S1 pocket making interactions with residues lining the S1 pocket as well as the catalytic triad. As a result, CDR-H3 paratope residues make several interactions with βFXIIa driving most of the affinity in contrast to CDR-H2 and CDR-L1. The residues in the CDR-H3 make a series of interactions with amino acids from the S1, S2 and S3 pocket of βFXIIa such as a “string of pearls” (**Fig. 2A**). Arg102 forms a network of hydrogen bonds with water molecules, side chain-main chain interactions with Val97. Arg102 is also oriented such that it sits on the top of the catalytic triad via a crucial stacking arrangement with Tyr99 and Trp215 of βFXIIa. Tyr105 sits tightly packed in the S1 pocket making main chain associations with Ala190, Ser195 and Gly216. Interestingly, Tyr105 also forms a hydrogen bond with Asp189 and is stacked between Ile213 and Trp215 in βFXIIa. Leu106 serves as a lid on the top of S1 pocket that Tyr105 occupies. The final “pearl of the string” is Tyr114 forming main chain interactions with Pro96 and Val97 (**Fig. 2A**). These interactions demonstrate mechanistically how CDR-H3 governs affinity towards βFXIIa and form the basis of βFXIIa inhibition.

The insertion of CDR-H3 into βFXIIa appears to govern tight binding and selectivity, with stabilizing interactions provided by CDR-H2 and CDR-L1. This is in line with the affinity maturation strategy of 3F7, wherein only GIRPSGGTTVYADSVKG residues in CDR-H2 were mutated to underlined residues GI<u>DIPTKG</u>TVYADSVKG. These changes resulted in a ∼44-fold increase in affinity. The introduction of a positively charged lysine in place of glycine at position 56 in CDR-H2 appears to increase main chain-main chain interactions as well as form new hydrogen bonds between the Arg60 side chain. Importantly, reorganization of the proline by mutation of Ser54 in the parental antibody has significantly matured the affinity through stacking interaction with Pro96 of βFXIIa (**Fig. 2A**). Comparison of 3F7-βFXIIa and garadacimab-βFXIIa binding kinetics data revealed that the interactions mediated by CDR-H2 have led to a slower dissociation rate, resulting in increased affinity (Cao et al., 2018). The high-resolution cryo-EM SPA of garadacimab Fab-βFXIIa complex also provides the basis for the potent inhibition of βFXIIa by garadacimab. Initial inspection of the overlay of uncomplexed βFXIIa (PDB id: 6gt6) (Pathak et al., 2019), benzamidine bound βFXIIa (PDB id: 6b74) (Dementiev et al., 2018) and garadacimab Fab bound βFXIIa did not show any significant conformational change in the overall structure (**Fig. S6**). Upon close inspection, it was evident that Tyr105 from CDR-H3 forms a salt bridge with Asp189 of βFXIIa in the same manner as the benzamidine (**Fig. 4B**). The mode of interaction is strikingly similar such that the phenyl ring of Tyr105 overlays exactly on the benzene ring of benzamidine. This points to a βFXIIa inhibition mechanism by garadacimab that is remarkably similar to the canonical inhibitor of serine proteases. The 99-loop in the benzamidine bound, and garadacimab complexed, βFXIIa appears to be in an identical conformation, wherein Tyr99 is in a closed form rendering the S2 pocket shut (**Fig. 4B**). This further demonstrates a conserved inhibition mechanism of βFXIIa. On the contrary, in the uncomplexed βFXIIa structure, this loop slightly moves away from the catalytic triad revealing an open form of Tyr99. Such a movement results in the opening of the S2 pocket and partial blockage of the S3 region. Altogether, this suggests that Tyr99 potentially serves as a switch for substrate binding.

The interactions mapped using the cryo-EM structure of Garadacimab Fab-βFXIIa were aligned against binding interfaces identified in our recent report using HDX-MS (Ow et al., 2023). Highlighted HDX-MS interfaces on βFXIIa show very good agreement with contact regions observed in the cryo-EM structure. Specifically, we demonstrate that the interfaces neighbouring the S1 catalytic pocket entrance (e.g. 60-loop, 99-loop, 180-loop and 220-loop) overlapped with regions shown to be blocked by the CDR-H3, CDR-H2 and CDR-L1 of garadacimab (Ow et al., 2023).

In conclusion, our study presents a high-resolution cryo-EM structure of garadacimab Fab-βFXIIa complex that most likely reflects how garadacimab engages with activated FXII. The structural analysis provides first insights into garadacimab governed inhibition mechanism of FXIIa. This appears to be in line with the downregulation of FXIIa proteolytic activity by its endogenous inhibitor, C1-INH.

## METHODS

### Protein production and purification

HuFXII (354-596)-epea was expressed in Expi293 expression system as per the manufacturers protocol. Briefly, Expi293 cells and the mammalian expression vector pcDNA3.1 were obtained from Thermo Fisher Scientific, USA. Transient transfections of expression plasmid encoding HuFXII (354-596)-epea were performed using ExpiFectamine™ according to the manufacturer’s recommendations (Thermo Fisher Scientific, USA). The cells were maintained at 37 °C and 8% CO_2_.

1 litre of culture supernatant was separated from the cells transiently expressing HuFXII (354-596)-epea. The supernatant was filtered with a 0.22 βM filter and loaded onto a tricorn 10/100 Capture Select C-tag XL™ resin (GE Healthcare, USA). The column was pre-equilibrated with 25 mM Tris, 2.7 mM KCl, 137 mM NaCl (TBS pH 7.4). Post loading, the column was washed with TBS pH 7.4 buffer to remove any unbound and non-specific proteins. The protein was eluted with 20mM Tris, 2M MgCl_2_ pH 7.4. Subsequently, the eluate was loaded on a Superdex 26/600 column pre-equilibrated with 10 mM HEPES pH 7.4, 150 mM NaCl (HBS pH 7.4). The eluate was concentrated using Amicon ultra centrifugal filters (MilliporeSigma, USA) with a MW cutoff of 3 kDa, sterile filtered, and stored at -80 °C to 3.1 mg/ml.

Garadacimab Fab was expressed in Expi293 cells using the above-mentioned method. The Fab was purified from 1 ltr of supernatant using cation exchange HiPrep SP Sepharose HP 16×x10 cm column. The pH of the supernatant was adjusted to pH 5 followed by filtration through a 0.22 μM filter. The column was equilibrated with 50 mM sodium acetate, pH 5 and the pH-adjusted supernatant was directly loaded onto this column. The protein was eluted with 50 mM sodium acetate, 1M NaCl, pH5. Subsequently, the eluate was loaded onto a gel filtration HiLoad Superdex200pg column pre-equilibrated with PBS pH 7.4. The eluted protein was concentrated using Amicon ultra centrifugal filters (MilliporeSigma, USA) with a MW cutoff of 3 kDa, sterile filtered, and stored at −80 °C to 3.6 mg/ml.

### Complex formation

βFXIIa-garadacimab Fab-VHH complex was prepared by forming the complex of βFXIIa-garadacimab Fab first. The complex was prepared by mixing 1.25-fold molar excess of βFXIIa with the Fab portion of garadacimab and incubated at room temperature for 20 minutes. The complex was isolated by size exclusion chromatography. To bulk up this complex, 1.5 molar excess VHH (Thermo Fisher Scientific (Catalog number: 7103082500)) was mixed with βFXIIa-garadacimab Fab complex to generate the ternary complex. The ternary complex was isolated and purified by size exclusion chromatography using a HiLoad 16/600 Superdex200 PG column in 10 mM Na Acetate, 100 mM NaCl pH 5.5. The relevant fractions were pooled using an Amicon Ultra-15 30kDa MWCO device. The complex was analysed using SDS-PAGE (4-12 % Novex precast Bis-TRIS gel) and size exclusion chromatography coupled with multi-angle static light scattering (SEC-MALS) using a dedicated S200 5/150 column at a 0.2 ml/min flow rate, 20-40 μg sample injected was used for final species and molecular weight analysis Analysis was performed in ASTRA 8.1.

### Cryo-EM sample preparation and data collection

Tertiary complex was diluted five times to the final concentration of 2 mg/ml. Immediately before blotting and plunge freezing, fluorinated octyl maltoside (FOM) was added to the sample to the final concentration of 0.005%-0.01% (w/v). 3 μl of the complex was applied to glow-discharged (20 mA, 30 s; Quorum GloQube) Quantifoil R1.2/1.3 grids (Quantifoil Micro Tools GmbH), blotted for 5 s using blot force 0 and plunge-frozen into liquid ethane using Vitrobot Mark IV (Thermo Fisher Scientific).

The sample was imaged on a Thermo Scientific Krios™ G4 Cryo-TEM operating at 300 keV. The microscope was equipped with an E-CFEG, a Thermo Scientific Selectris™ X Energy Filter, and a Falcon 4i Detector operated in Electron-Event Representation (EER) mode. EPU 2 automated acquisition software was used to collect 4000 movies (2000 with 0.005% FOM and 2000 with 0.01% FOM) at 0.73 Å/pixel with a total dose of 50 e^-^/Å^2^. Defocus targets cycled from -0.5 to -1.25 microns.

### Cryo-EM data processing

Data processing was performed using the cryoSPARC software (Punjani et al., 2017). After patch-motion and patch CTF correction, particles were picked from each dataset independently using a blob picker, extracted and subjected to 2D classification to generate templates for the template picker. Following template picking, particles were extracted in a 280-pixel box (pixel size 1.09 Å/px) and 2D classified. Particles belonging to 2D class averages with high-resolution features were selected for Ab initio reconstruction into three classes. Following independent Ab Initio reconstructions for each of the datasets, ab initio classes showing similar architecture were combined and subjected to non-uniform refinement (Punjani et al., 2020), resulting in 2.6 Å and 3.6 Å reconstructions for the open and closed dimer, respectively. Due to the low occupancy of one of the protomers in the closed dimer, 3DVAR analysis (Punjani & Fleet, 2021) was performed using the particles in the closed dimer reconstruction using a global mask. Clusters were combined depending on whether they displayed a dimer or a monomer and refined separately to produce 3.8 and 3.9 Å reconstructions (C1), respectively. Directional FSC was estimated using 3DFSC (Aiyer et al., 2021). Data processing workflow and relevant information are presented in Figure S4 and Table S1.

### Model building, refinement, and interpretation

UCSF Chimera X (version 1.4) and Coot (version 1.0) were used for model building and analysis (Emsley & Cowtan, 2004; Pettersen et al., 2021). The crystal structure of βFXIIa (PDB id: 6b74) (Dementiev et al., 2018) was fitted into our density using the UCSF Chimera X “Fit in map” tool. Fab atomic model was predicted from sequence information using Phyre2 (Kelley et al., 2015). CDR regions were built manually in the density map using Coot (Emsley & Cowtan, 2004). The models were refined against the respective EM density maps using Phenix Real Space Refinement and validated with MolProbity (Liebschner et al., 2019; Williams et al., 2018). PDBsum was used to identify βFXIIa residues interacting with garadacimab (Laskowski et al., 2018). Figures were generated using UCSF ChimeraX (Pettersen et al., 2021).

## Data availability

Coordinated of the garadacimab Fab-βFXIIa complex are deposited in the Protein Data Bank under the accession code 8R8D. The EM density maps of the C2 dimer, C1 dimer and C1 monomer have been deposited to the Electron Microscopy Data Bank under the accessions EMD-18999, EMD-19000 and EMD-19001, respectively. All data needed to evaluate the conclusions in the paper are present in the paper and/or the Supplementary Materials. Additional data related to this paper may be requested from the authors.

## Acknowledgements

We thank CSL R&D (Marburg, Germany) for providing the purified βFXIIa material and the CSL R&D Lab Operations Team (Melbourne, Australia) for organising the transport and shipping of materials to Thermo Fisher Scientific for cryo-EM data collection (Center for Electron Microscopy, Eindhoven NanoPort, The Netherlands). We are grateful to Boyin Liu, Mazdak Radjainia, Kasim Sader, and Ryan Shaw (Thermo Fisher Scientific) for their support, knowledge sharing and management of this project.

## Contributions

M.P., C.P., E.A.K conceived the project. I.D., R.G., A.J.Q., E.A.K., M.P. designed the experiments. I.D., M.P., R.G., A.J.Q., A.D.N., M.J.W. provided access to equipment and reagents. A.J.Q., R.G. prepared protein complexes. I.D. carried out cryo-EM sample preparation, data collection, and analysis. I.D., R.G., S.Y.O., M.P. analyzed, visualized the data and prepared figures. M.P. supervised the project. I.D., R.G., S.Y.O., M.P. wrote the first draft of the manuscript. All authors contributed to reviewing and editing subsequent versions.

