## supplemental for "Structural basis for inhibition of βFXIIa by garadacimab"

**This PDF file includes:**

Figs. S1 to S8

Table S1

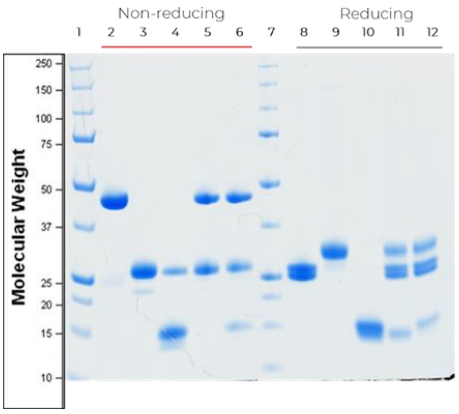

| Lane | Sample | Vol (ul) |
|---|---|---|
| 1 | BioRad Precision Plus Mw markers | 10 |
| 2 | 3F7 VR115 Fab (Non-reducing) | 1 |
| 3 | FXIIa beta C467S (Non-reducing) | 1 |
| 4 | Biotin-anti-lambda VHH (Non-reducing) | 3 |
| 5 | 3F7 VR115 Fab-FXIIa beta-VHH (HMW) (Non-reducing) | 0.5 |
| 6 | 3F7 VR115 Fab-FXIIa beta-VHH (Non-reducing) | 0.3 |
| 7 | BioRad Precision Plus Mw markers | 10 |
| 8 | 3F7 VR115 Fab (Reducing) | 1 |
| 9 | FXIIa beta C467S (Reducing) | 1 |
| 10 | Biotin-anti-lambda VHH (Reducing) | 3 |
| 11 | 3F7 VR115 Fab-FXIIa beta-VHH (HMW) (Reducing) | 0.5 |
| 12 | 3F7 VR115 Fab-FXIIa beta-VHH (Reducing) | 0.3 |

**Figure S1:** SDS-PAGE analysis of the individual, binary and ternary complex components of the pooled gel filtration fractions.

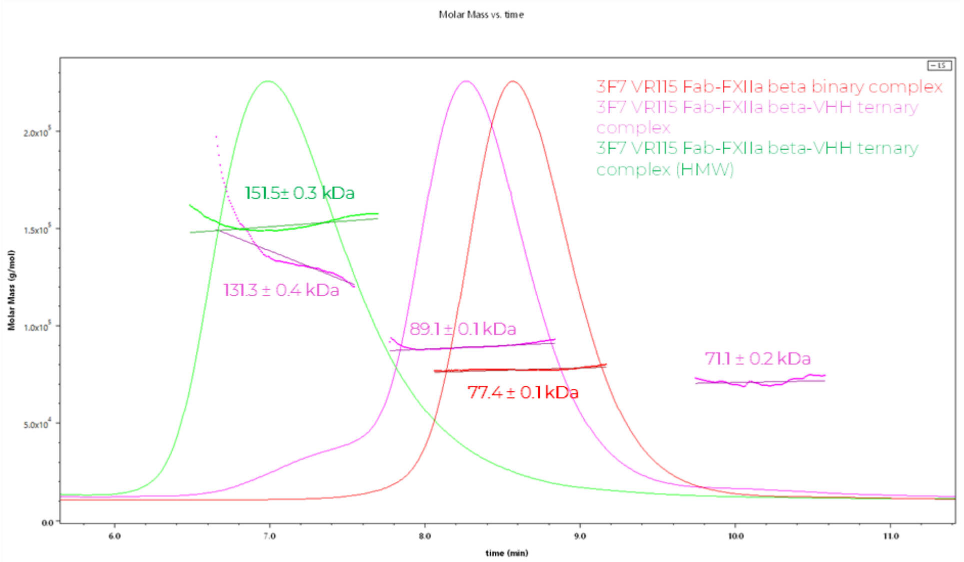

**Figure S2:** SEC-MALS analysis of the binary (red), ternary (pink) and high-molecular-weight ternary (green) complex components.

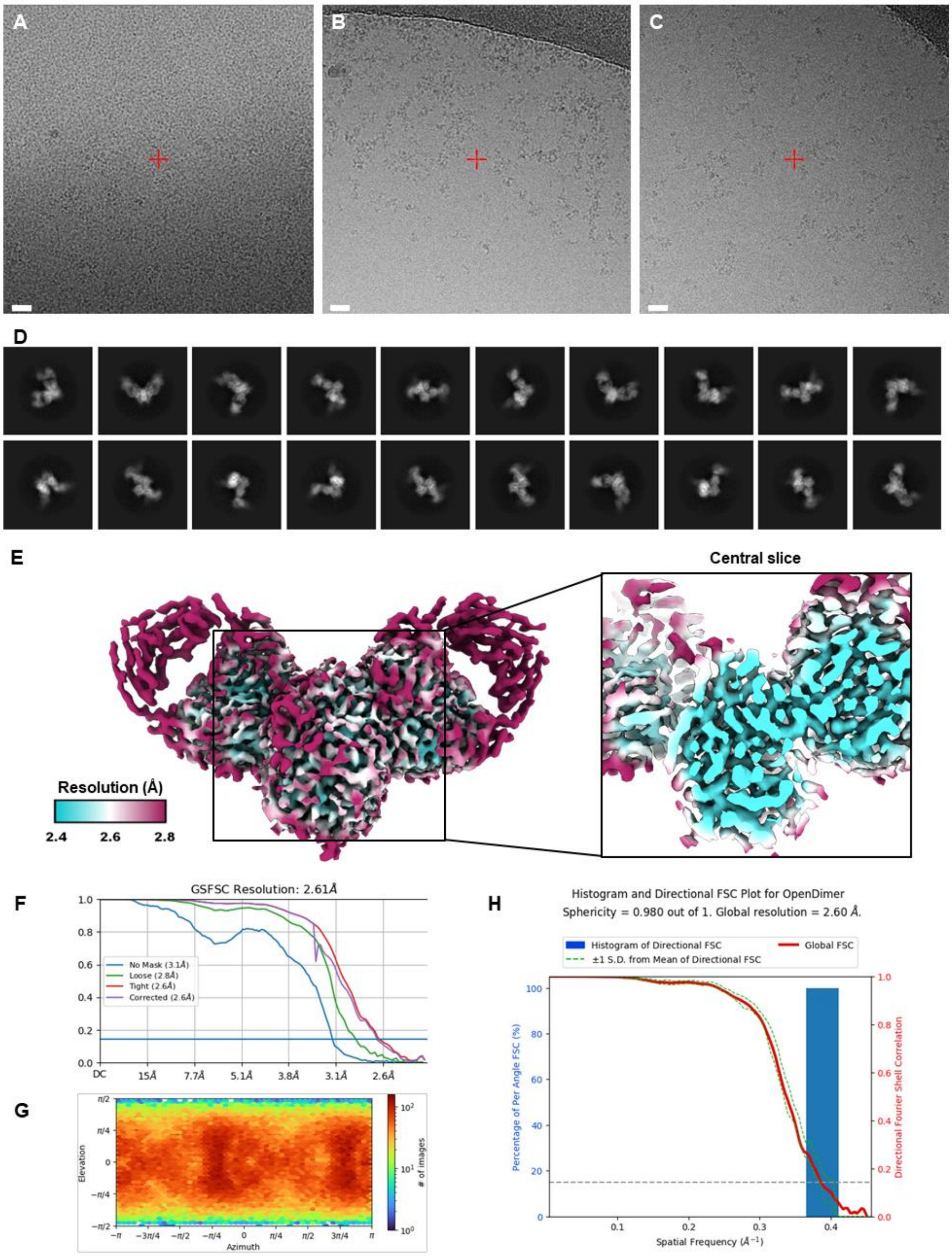

**Figure S3: Cryo-EM analysis of the C2-symmetric garadacimab fab-βFXIIa complex.** Raw micrographs of the ternary complex (A) without fluorinated octyl maltoside, and with (B) 0.005% or (C) 0.01% FOM. Scale bar = 20 nm. (D) Representative 2D class averages of the C2-symmetric garadacimab fab-βFXIIa dimer. (E) Local resolution filtered EM density map for the C2 refined garadacimab fab-βFXIIa dimer, coloured according to local resolution, which was calculated in cryoSPARC. An inset shows a central slice of the paratope-epitope region, resolved to ~2.4 Å. (F) Gold-standard Fourier shell correlation (FSC) curve generated from the independent half maps contributing to the 2.6 Å global resolution density map of the garadacimab fab-βFXIIa complex. (G) Angular distribution plot and (H) 3DFSC plot for the C2 refined garadacimab fab-βFXIIa dimer.

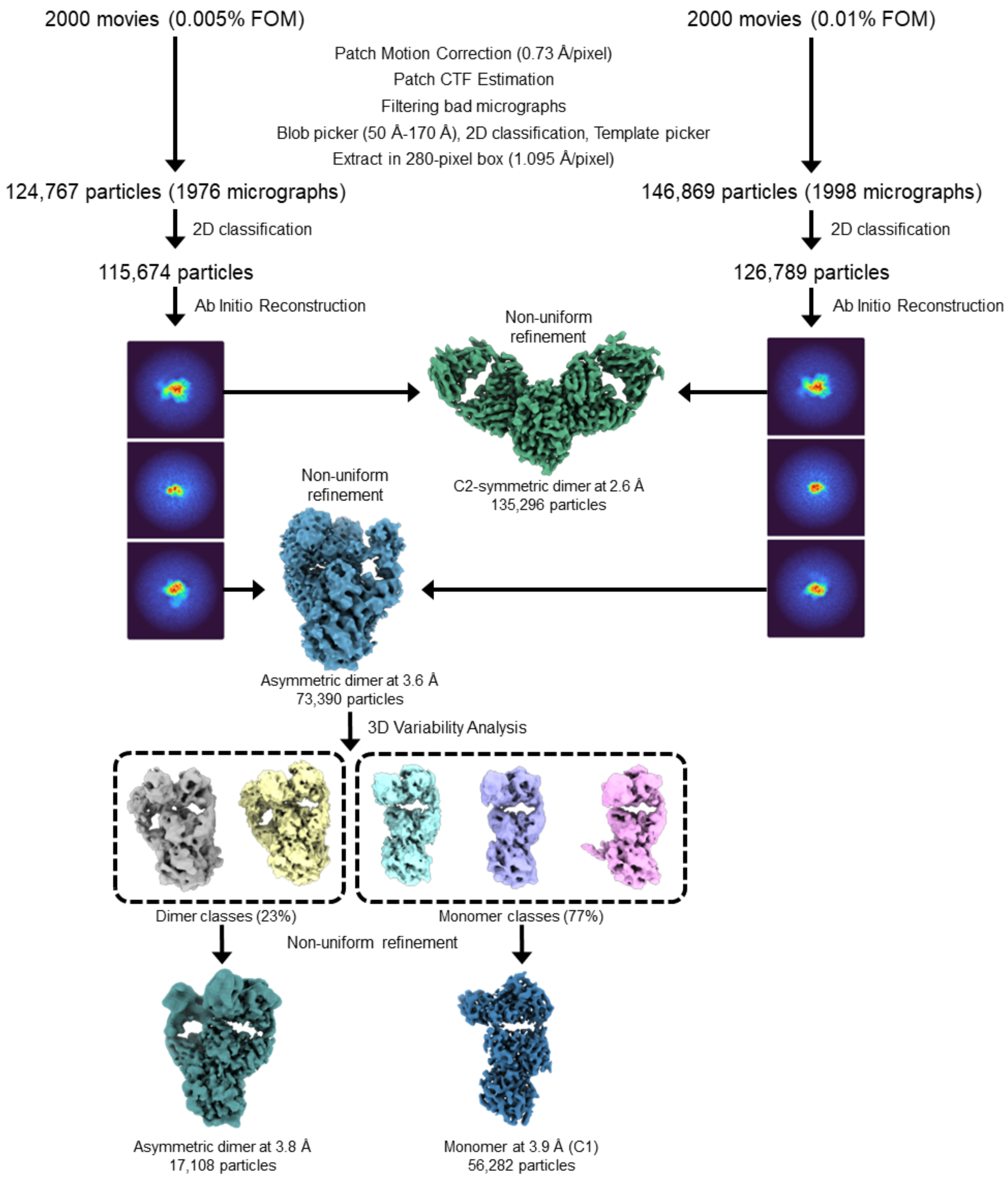

**Figure S4:** Single-particle cryo-EM image processing workflow for the garadacimab fab-βFXIIa complex.

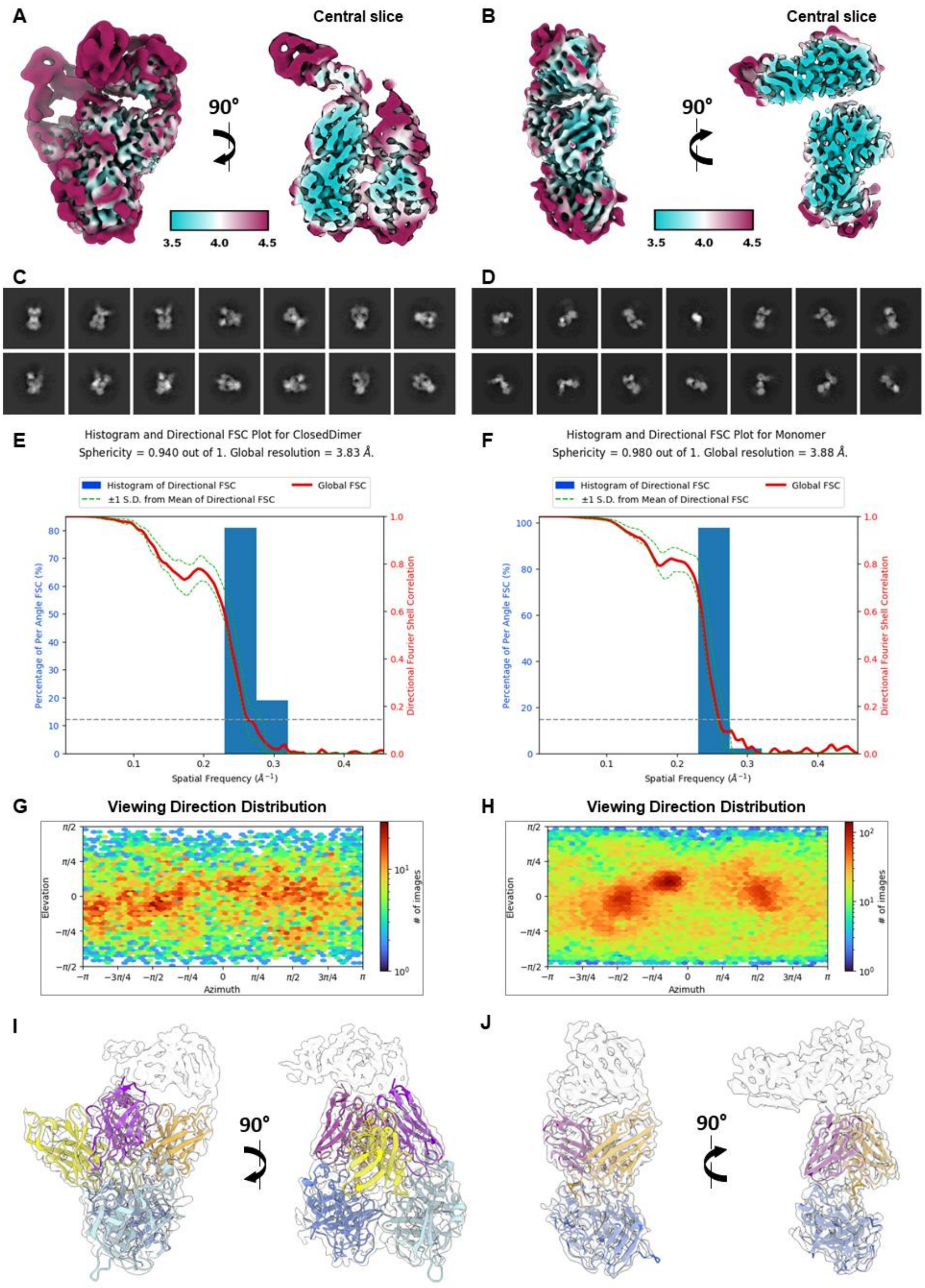

**Figure S5: Cryo-EM analysis of the alternative garadacimab fab-βFXIIa complex conformation.** Local resolution filtered EM density map for the C1 refined garadacimab fab-βFXIIa dimer (A) and monomer (B), coloured according to local resolution, and shown as two orthogonal views. Representative 2D class averages of the C1 garadacimab fab-βFXIIa dimer (C) and monomer (D). Histogram and Directional Fourier Shell Correlation Plots for the C1 refined garadacimab fab-βFXIIa dimer (E) and monomer (F). Angular distribution plots of the C1 refined garadacimab fab-βFXIIa dimer (G) and monomer (H) maps. Transparent EM density map of the C1 refined dimer (I) and monomer (J) showing the rigid-body fitted atomic model of garadacimab fab-βFXIIa complex.

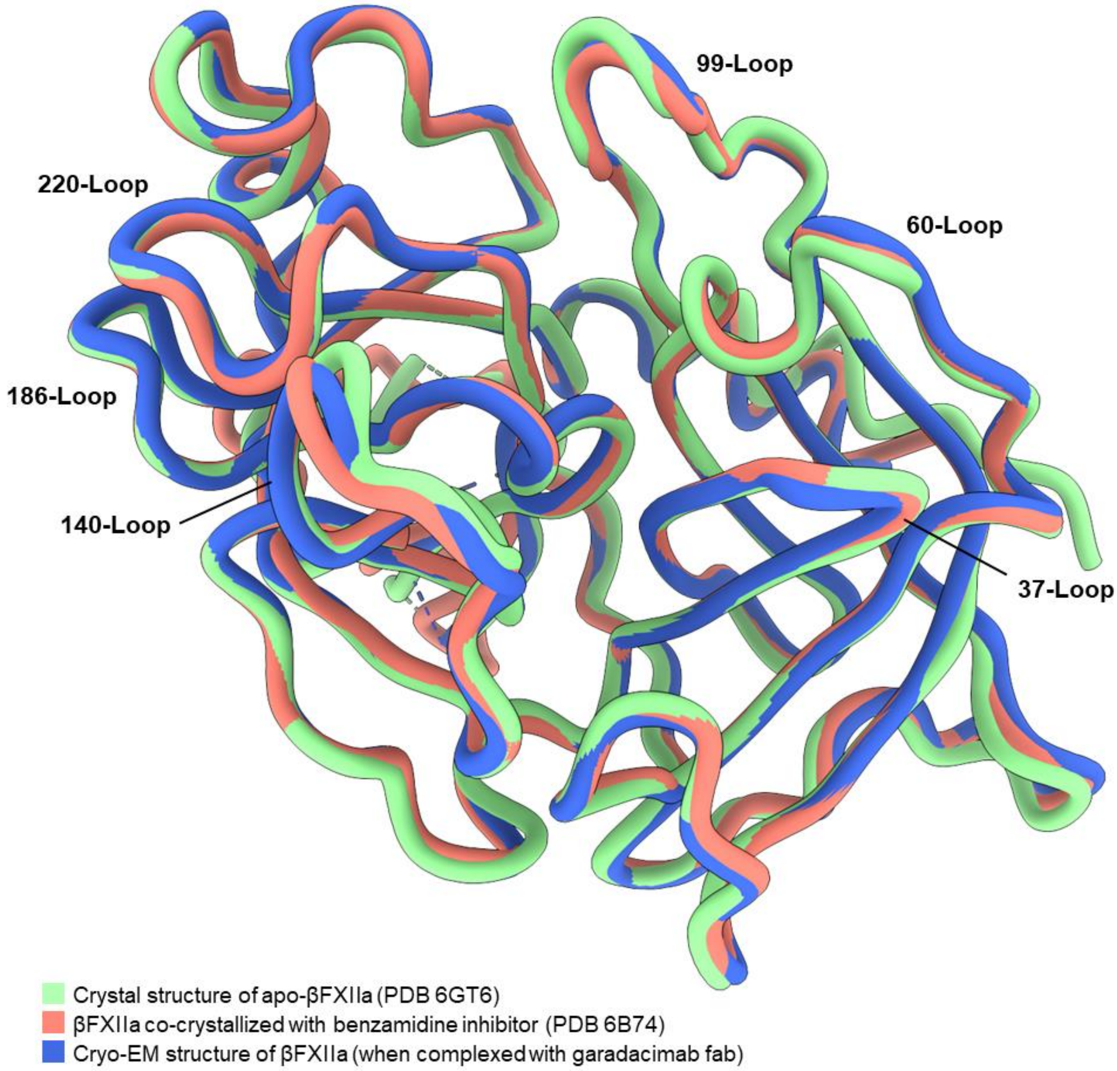

**Figure S6:** Cryo-EM structure of Factor XIIa (blue) shows little differences when overlaid with the crystal structures of βFXIIa apo (green) and holo (red).

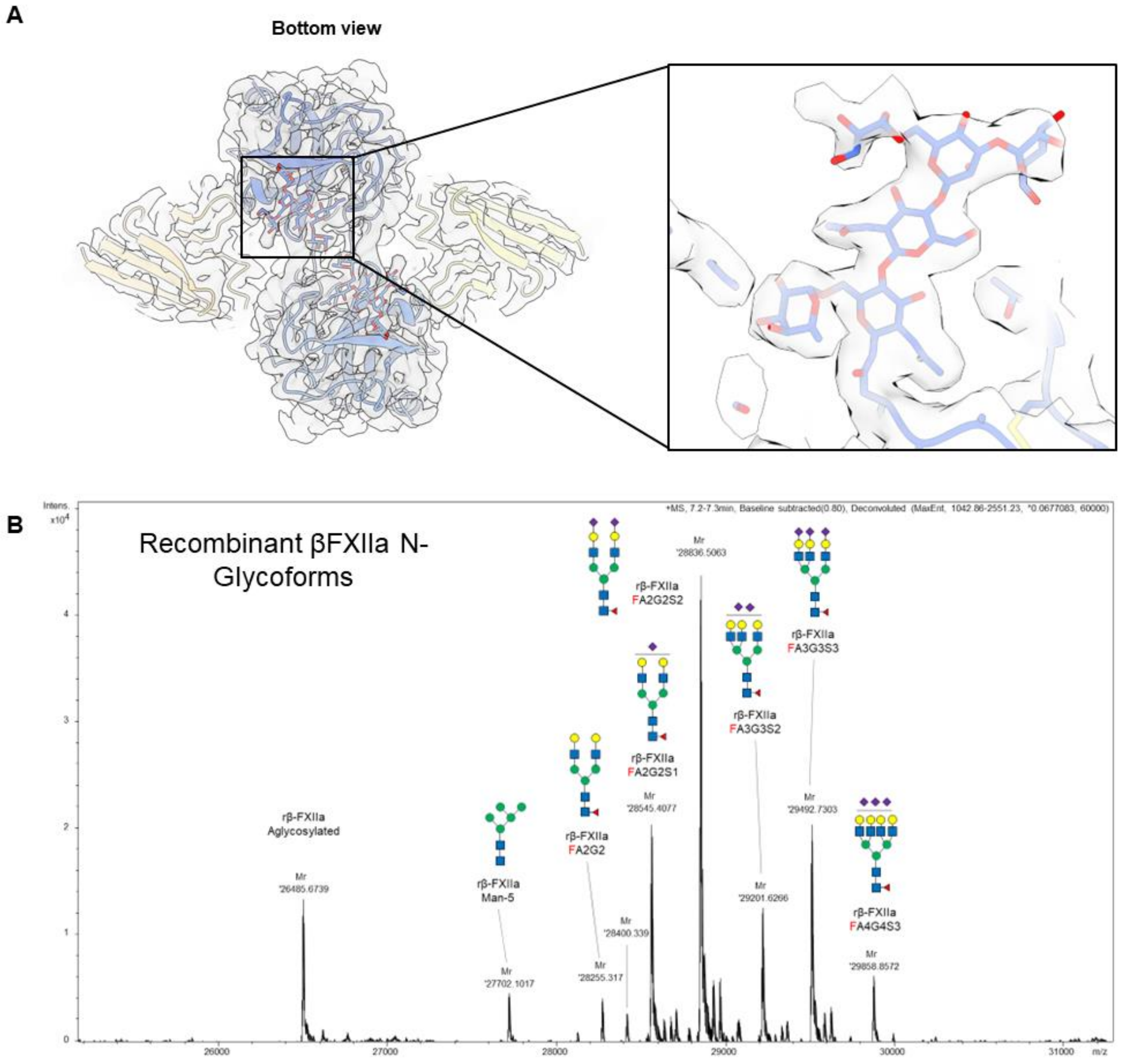

**Figure S7:** (A) Sialylated bi-antennary core-focusylated glycan density near the βFXIIa dimerization interface. (B) Mass spectrum confirming the identity of the observed glycan. Molecular mass analysis of HuFXII(354-596)-epea [C467S] was performed using a high-performance reverse phase liquid chromatography coupled with high resolution time-of-flight mass spectrometer. Briefly, two micrograms of protein sample were injected onto a MAbPac RP 2.1×50mm, 4 µm, 1500Å high performance reverse phase column (ThermoFisher, Waltham, MA US). Protein samples were held at 300 µL/min for a 2-minute desalting step, followed by a 6-minutes 5-50% linear gradient consisting of 0.1% formic acid in Milli-Q water against 0.1% formic acid in acetonitrile at the same flowrate. The method includes a further 7-minute washing and conditioning, with a total run time of 15 minutes. The separated proteins were detected using an electrospray coupled maXis 4G Qq-TOF mass spectrometer (Bruker Daltonik, Bremen DE). Mass spectrometer capillary voltage was set at 4 kV with an endplate offset of 500V. Electrospray source and dry gas at 0.8 bar and 0.8 L/min respectively. Ion transmission settings were set to manufacturer’s default for intact proteins. Mass spectra were recorded at 0.5 scans per second at a mass range of 900-5000 m/z.

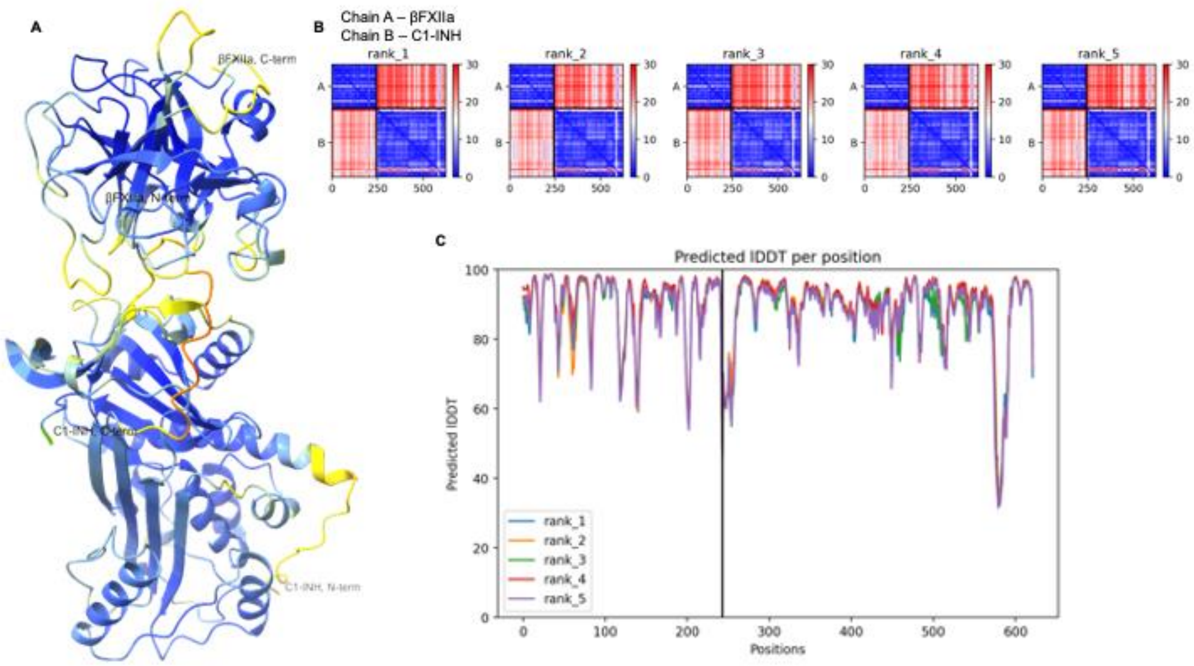

**Figure S8:** (A) Cartoon of alphafold generated model of C1-INH in complex with βFXIIa colored as per pLDDT (predicted local distance difference test) values. (B) PAE plots of all the five models of C1-INH in complex with βFXIIa. (C) Plot demonstrating predicted local Distance Difference Test (LDDT) per position for the five models.

**Table S1:** Cryo-EM data collection, refinement and validation statistics.

| | #1 Garadacimab Fab-βFXIIa C2-symmetric dimer (EMD-18999) (PDB 8R8D) | #2 Garadacimab Fab-βFXIIa asymmetric dimer (EMD-19000) | #3 Garadacimab Fab-βFXIIa monomer (EMD-19001) |
|---|---|---|---|
| **Data collection and processing** | | | |
| Magnification | 165,000 | 165,000 | 165,000 |
| Voltage (kV) | 300 | 300 | 300 |
| Electron exposure (e–/Å$^2$) | 50.0 | 50.0 | 50.0 |
| Defocus range (μm) | -0.5 to -1.25 | -0.5 to -1.25 | -0.5 to -1.25 |
| Pixel size (Å) | 0.73 | 0.73 | 0.73 |
| Symmetry imposed | C2 | C1 | C1 |
| Initial particle images (no.) | 271,636 | 271,636 | 271,636 |
| Final particle images (no.) | 135,296 | 17,108 | 56,282 |
| Map resolution (Å) | 2.6 | 3.8 | 3.9 |
| FSC threshold | 0.143 | 0.143 | 0.143 |
| Map resolution range (Å) | 2.45-37.3 | 3.3-56.9 | 3.35-63.0 |
| FSC threshold | 0.5 | 0.5 | 0.5 |
| **Refinement** | | | |
| Initial model used (PDB code) | 6b74.pdb | | |
| Model resolution (Å) | 2.5 Å | | |
| FSC threshold | 0.143 | | |
| Model resolution range (Å) | 2.3-2.7 | | |
| Map sharpening *B* factor (Å$^2$) | -82.6 | -65.7 | -109.8 |
| Model composition | | | |
| Non-hydrogen atoms | 7118 | | |
| Protein residues | 928 | | |
| Ligands | BMA: 2<br>NAG: 4<br>FUC: 2<br>MAN: 4 | | |
| *B* factors (Å$^2$) | | | |
| Protein (min/max/mean) | 3.15/49.26/14.77 | | |
| Ligand (min/max/mean) | 11.28/38.76/22.7 | | |
| Water (min/max/mean) | 3.16/7.20/4.81 | | |
| R.m.s. deviations | | | |
| Bond lengths (Å) | 0.003 | | |
| Bond angles (°) | 0.581 | | |
| Validation | | | |
| MolProbity score | 1.68 | | |
| Clashscore | 6.70 | | |
| Poor rotamers (%) | 0 | | |
| Ramachandran plot | | | |
| Favored (%) | 95.67 | | |
| Allowed (%) | 4.52 | | |
| Disallowed (%) | 0 | | |
